# Molecular and biological characterization of a distinct species of *Lolavirus* infecting different accessions of seashore paspalum, a turfgrass, widely grown in the United States

**DOI:** 10.1101/2025.03.07.642087

**Authors:** Sayanta Bera, Taylor F. Schulden, Xiaojun Hu, Peter Abrahamian, Yu Yang, Anna L. Paulson, Amy Harvey-White, Shreena Pradhan, Katrien Devos, Christina Devorshak, Joseph A. Foster, Bishwo N. Adhikari

## Abstract

Seashore paspalum (*Paspalum sp*.), is an economically significant grass used in golf courses, sports fields, and landscaping in the United States. A novel *Lolavirus*, tentatively named paspalum latent virus (PaLV), was identified for the first time in seashore paspalum plants from the USDA National Plant Germplasm System (NPGS) using high-throughput sequencing. Three complete genome sequences of PaLV from different *Paspalum* accessions, with a length of 6,995 nucleotides (nt), not including the poly(A) tail, were obtained by Rapid Amplification of cDNA Ends and Sanger sequencing. Phylogenetic analysis based on the replicase protein sequences from the *Alphaflexiviridae* family revealed that PaLV grouped with the *Lolavirus* genus, with the closest relative being Lolium latent virus (LoLV). PaLV shares less than 72% nt identity to the replicase and coat protein genes of LoLV, which demarks PaLV as a new species and the second member of the genus. Furthermore, the coat protein region showed intense negative selection pressure and low spatially structured diversity. Host range analysis of PaLV showed that wheat, corn, sorghum, and *Lolium* are systemic hosts of PaLV. A one-step RT-PCR technique was developed to reliably detect PaLV infection.

## Introduction

Turfgrasses play a vital role in environmental protection and human well-being^1^. Seashore paspalum (*Paspalum vaginatum* Sw.), a warm-season perennial turfgrass, has attracted attention due to its salinity tolerance and adaptability to coastal environments in tropical and subtropical regions ^2,3^. As a result, seashore paspalum is a popular choice for sports fields in coastal regions where freshwater availability is an issue and saltwater irrigation results in tremendous cost savings ^4^. Beyond its functional benefits, seashore paspalum also contributes to environmental sustainability by mitigating the urban heat island effect, providing forage and aesthetic value, and generating economic and social benefits. The species is also reportedly tolerant to flooding and low oxygen conditions ^1,5,6^. Due to these numerous benefits, the cultivation area of *Paspalum* has expanded beyond coastal areas; however, in new environments, plants are exposed to many unknown pathogens, increasing the risk of disease damage.

Seashore paspalum, similar to other turfgrasses, is propagated through clonal methods, resulting in genetic uniformity ^7^. As a result, viruses are easily spread across the propagated plant material and introduced into new areas ^7,8^. Notable viruses that have been identified in *Paspalum*, under natural or experimental conditions, include *Paspalum striate mosaic virus* (PSMV, *Geminiviridae*) ^9,10^, *Sugarcane mosaic virus* (SCMV, *Potyviridae*) ^11^, *Sugarcane streak Réunion virus* (SSRV, *Geminiviridae*) ^12^, *Chloris striate mosaic virus* (CSMV, *Geminiviridae*) ^9^, *Paspalum dilatatum striate mosaic virus* (PDSMV, *Geminiviridae*) ^9^, and *Barley yellow dwarf virus* (BYDV, *Tombusviridae*) ^13^.

In recent years, high-throughput sequencing (HTS) has emerged as a transformative technology in the field of virology, particularly in the realm of viral discovery and diagnostics ^14,15^. This approach has led to the identification of numerous novel viruses in diverse biological samples, including grasses, weeds, trees, and food crops ^16–20^. Therefore, in this study we leveraged HTS to investigate the virome of *Paspalum* germplasm used in breeding programs. For this purpose, 27 *P. vaginatum* and three *P. distichum* (a sister species of *P. vaginatum*) accessions were obtained from the USDA National Plant Germplasm System (NPGS) collection and maintained at the University of Georgia (UGA), Griffin and Athens campuses.

Our results include the first report of a *Lolavirus* infecting both *P. vaginatum* and *P. distichum*. Confirmation of putative viral sequences was obtained by RT-PCR. The complete viral genome was obtained by 5’ RACE, indicating a novel species according to the species demarcation criteria of the genus *Lolavirus* ^21^, tentatively named Paspalum latent virus (PaLV). Here, we report the identification of PaLV, its biological and molecular characterization, phylogenetic relationships, genetic diversity and population structure, and PaLV screening of *P. vaginatum and P. distichum* germplasm maintained in greenhouses at UGA.

## Material and Methods

### Plant Material and Growth

*Paspalum* plants, obtained initially from the USDA-NPGS collection, Griffin, were maintained in greenhouses at the UGA Griffin and Athens campuses in 4×4 pots. The soil was a 1:1 mixture of sand and Miracle-Gro potting mix. Plants were fertilized once every two weeks with Osmocote Plus.

### Total Plant RNA Extraction

Total RNA from the *Paspalum* accessions PI 509022 leaf tissue was extracted using a RNeasy Plant Mini Kit (Qiagen, Hilden, Germany), and an RNase-Free DNase Set (Qiagen) was used for DNA digestion. The manufacturer’s recommended protocol was used without modifications. For accessions HI10 and Spence, papillae were peeled from the adaxial surface of leaves (second fully open, developed leaf on growing stolon segment) from plants grown under freshwater (0 mM NaCl) and salt stress (200 mM NaCl) for six weeks. Strings of papillae were collected in 2 mL tubes resting on a liquid nitrogen bath to maintain RNA integrity, and frozen tissues were ground using a TissueLyser II bead mill (Qiagen). The ground tissue was homogenized in 1 mL of TRIzol reagent (Invitrogen, Waltham, MA) per 100 mg of tissue, followed by phase separation after adding 200 μL of chloroform. The RNA-containing aqueous phase was carefully transferred to a 2 mL nuclease-free tube and mixed with an equal volume of 100% ethanol. The resulting RNA-ethanol mixture was loaded onto a Zymo-Spin IC column from the Zymo RNA Clean & Concentrator-5 kit (Zymo Research, Irvine, CA), and RNA clean-up was performed following the manufacturer’s instructions. RNA was eluted in 10 μL nuclease-free water and RNA concentration was determined using a NanoDrop spectrophotometer, and the quality was assessed via electrophoresis on a 1% Tris-Borate-EDTA (TBE) agarose gel.

### HTS and Virus Identification

The RNA extracts from leaves of accession PI 509022 at USDA APHIS-PPQ, Beltsville, and from adaxial leaf papillae from accessions HI 10 and Spence at the University of Georgia, Athens, were subjected to HTS at their respective place of extraction. For HTS of RNA from PI 509022, a single-indexed ribosomal-RNA-depleted cDNA library was prepared using the TruSeq® Stranded Total RNA Library Prep Plant kit (Illumina, San Diego, CA) and sequenced on an Illumina NextSeq 500 platform, generating 19,815,890 single-end, 75-bp reads. For HTS of RNA from accessions HI 10 and Spence, barcoded stranded mRNA-seq libraries were generated from 500 ng of RNA with the KAPA Stranded mRNA-Seq Kit (Roche, Basel, Switzerland) according to the manufacturer’s instructions, using half-reactions for all steps except the final library amplification. The concentration of the libraries was measured using a Qubit Fluorometer with the Qubit 1X dsDNA High Sensitivity Assay Kit (Invitrogen). Subsequently, 30 ng of each library was pooled into a larger set of 24 leaf and peels libraries and loaded onto a single flow cell of an Illumina NextSeq 2000 for paired end (PE) 150 bp sequencing at the Georgia Genomics and Bioinformatics Core (GGBC) at UGA. Reads from HTS were assembled using the de novo assembly tool in PhytoPipe (2023) ^22^ and compared with viral pathogen databases. Briefly, raw reads were filtered and trimmed using Trimmomatic (v.0.39) ^23^. Then, contigs were assembled using Trinity (v2.8.6) ^24^ and compared to the NCBI Viral Reference Database (Jan 2022) by blastn search ^25^ and to the Reference Viral Database protein database (v22)^26^ by Diamond blastx search ^27^. Default parameters were used in all programs.

### 5’ RACE

Rapid amplification of cDNA ends with the SMARTer® RACE 5′/3′ Kit (Takara) was used to complete the 5′ end of the ssRNA segment with the RACE primer – 5’**-** GGACCACTGTGGCGTAGAGGATGTCGAGGT-3’; the 5’ end of the primer was linked to the adapter sequence provided by the company. All PCR fragments were sequenced by direct Sanger sequencing in both directions, and sequences were aligned to the reference HTS-derived contigs using Geneious (v11.0.3).

### RT-PCR Validation of HTS and Screening of Germplasm

A one-step RT-PCR protocol was developed for detection of PaLV using the SuperScript™ III One-Step RT-PCR System with Platinum® Taq DNA Polymerase (Invitrogen). Primer pairs ORF1.FP: 5′-AAACAAGTGGAATTTCACGGG-3′ and ORF1.RP: 5′-CTGCTCTGGTGAGAAGATATCG-3′, were designed to amplify a 605 bp amplicon from the replicase gene. Cycling parameters were used following the manufacturer’s guidelines except for the 62°C annealing temperature.

### Host Range Studies

After confirmation of PaLV infection in *Paspalum* through RT-PCR, infected plant tissues were freshly ground in 0.01M Phosphate buffer (pH 7) for mechanical inoculation. All plants were sprayed with carborundum, an abrasive, before rub-inoculation. Mechanical inoculation of sap was conducted on nine different plant species: *Hordeum vulgare, Triticum aestivum, Zea mays, Sorghum spp*., *Dactylis glomerata, Setaria italica, Lolium multiflorum, Miscanthus sacchariflorus*, and *Avena sativa*. Three biological replicates were used for each species. All bioassay plants were kept in controlled-temperature greenhouse conditions at 26□ ± 2□ with 16 h light and 8 h dark. Two plant species, *H. vulgare*, and *T. aestivum* were assayed in a growth chamber maintained at 16□ with 16 h light and 8 h dark.

### Sequence, Phylogenetic, and Recombination Analysis

The viral sequences were analyzed by blastn (https://blast.ncbi.nlm.nih.gov/Blast.cgi) with default parameters. The putative proteins and potential open reading frames (ORFs) were determined using ORFfinder (https://www.ncbi.nlm.nih.gov/orffinder/) and subsequent blastp annotation by using the non-redundant protein sequences database. Pairwise comparisons between related viral ORFs were performed with Sequence Demarcation Tool (SDT) v1.2 to calculate the identity percentage for each ORF^28^. Conserved domains within these proteins were identified using the Conserved Domain Database (CDD) ^29^. The coat protein (CP) sequences were modeled using AlphaFold2^30^. PaLV CP (CP_PaLV_) was superimposed on the Lolium latent virus (CP_LoLV_) using TM-align ^31^. Structural similarity between CP_PaLV_ and CP_LoLV_ was further evaluated based on TM scores, using 0.5 as a cut-off. Superimposed images were exported from TM-align and loaded into Chimera X ^32^ for further processing.

Evolutionary relationship of PaLV with representative species from the genera, *Lolavirus, Potexvirus, Mandarivirus*, and *Allexivirus* in the *Alphaflexiviridae* family were included in a maximum-likelihood phylogenetic tree based on amino acid sequences of replicase using the LG with gamma distribution and frequency substitution model previously inferred using jModelTest in MEGA 11 ^33^. The phylogenic analysis was performed with 1,000 bootstrap replicates and potato virus T as an outgroup. The resulting phylogenetic tree was visualized and formatted using FigTree v1.4.4 (http://tree.bio.ed.ac.uk/software/figtree/).

The Recombination Detection Program V.5 (RDP5) program was used to explore the occurrence of recombination events in full-length viral genome sequences. Various methods implemented in RDP5 ^34^ including RDP, SisterScan, Bootscan, Chimaera, GeneConv, MaxChi, and 3Seq algorithms, were used. Any recombinant events with *P* < 0.001 predicted by four of the seven prediction methods were considered reliable and used in the analysis.

### Genetic diversity

The genetic diversity of the PaLV population was analyzed using 11 PaLV isolates obtained from different *Paspalum* accessions from the UGA collection. Genetic diversity was calculated based on the complete CP gene nucleotide sequences. The primer set designed to amplify the CP region of PaLV was developed according to the consensus sequence of three PaLV isolates obtained through HTS (CP_F: 5′-GATCGGAAGCCTCAGTTGTG-3′; CP_R: 5′-CGGTGGCCAGGGTAGATTAA-3′).

The CP nucleotide sequence of 11 PaLV isolates was aligned using the Muscle program implemented in MEGA 11 and the mean genetic diversity for the entire population was calculated. The average number of non-synonymous (d_N_) and synonymous (d_S_) nucleotide substitutions per site and the dN/dS ratio as an estimate of the CP’s selection pressure were computed using the DnaSP v.5 software ^35^. The Wright’s fixation index statistic *F*_*ST*_ ^36^ was calculated using DnaSP v.5. Frequent gene flow is considered to occur when *F*_*ST*_ < 0.33.

## Results and Discussion

### Genome structure

High-throughput sequencing identified three nearly complete genome sequences of PaLV infecting the *Paspalum* germplasm. The 5′ RACE was performed to determine the complete sequence of PaLV which consists of 6,995 nucleotides (nt) excluding the poly(A) tail, encodes five open reading frames (ORF) flanked by an 81 nt 5′ untranslated region (UTR) and 86 nt 3′ UTR. A pairwise comparison of each ORF of PaLV with ORFs from representative viruses revealed the highest degree of identity with LoLV, ranging from 49 to 57% at the nt level and 26 to 55% at the amino acid level (Table 1).

**Table 1:**
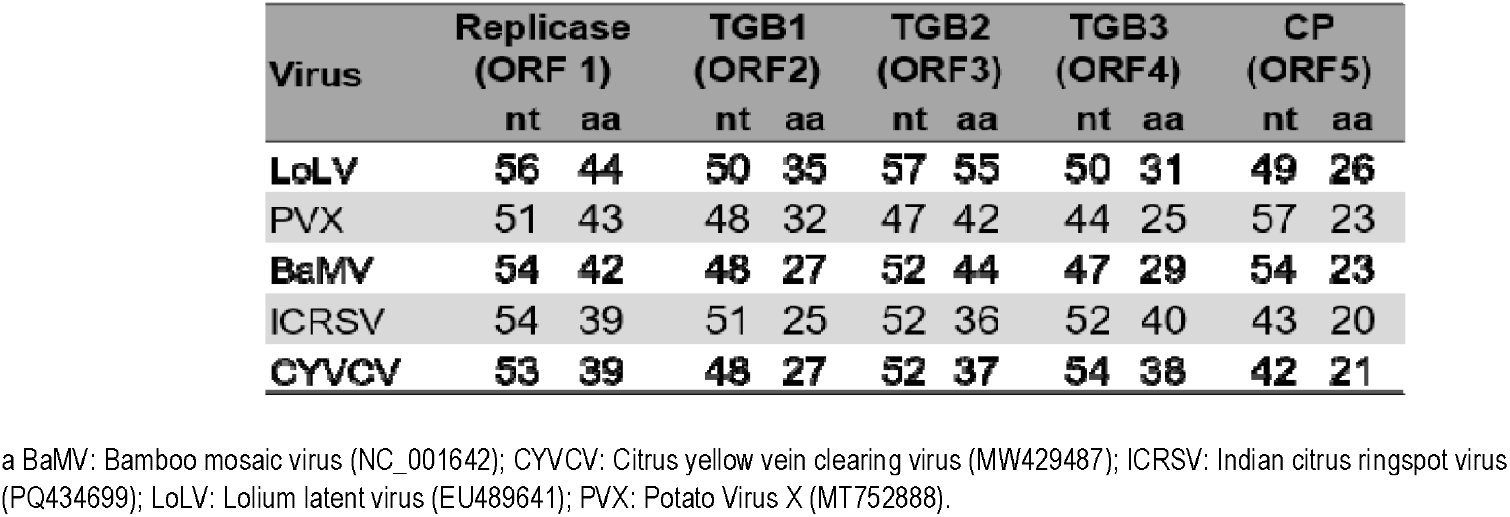
Identity (%) in nucleotide and deduced amino acid (aa) sequences between PaLV and other viruses from *Lolavirus, Potexvirus*, and *Mandarivirus* genera.

ORF1 is predicted to encode a 173-kDa protein of 1,524 amino acids (aa). A blastp analysis revealed maximum identity (56% and 44% in nt and aa, respectively) to ORF1 (replicase) of LoLV from the *Lolavirus* genus, *Alphaflexiviridae* family (Table 1). An exhaustive analysis of the amino acid sequence of ORF1 in CDD ^29^ revealed the presence of the following domains: Methyl transferase (aa 39-329 [7.90e-63]), Viral RNA helicase (aa 768-997 [1.04e-50]), and RNA-dependent RNA polymerase (aa 1178-1474[1.93e-167]) (Fig. 1A). However, the AlkB domain, which is present in LoLV, was absent in ORF 1 of PaLV. This is a notable characteristic of *Alphaflexiviridae*, where AlkB is present in some, but not all, viruses within the same genus. For instance, in the genus *Potexvirus*, papaya mosaic virus (PapMV) possesses AlkB, whereas potato virus X (PVX) and white clover mosaic virus do not ^37^. The AlkB domain is speculated to play a crucial role in maintaining the stability of RNA genomes, which are susceptible to methylation due to pesticide applications ^37,38^. This speculation led to the hypothesis that the acquisition of the AlkB domain is a relatively recent development in the evolution of viruses within the *Alphaflexiviridae* family ^37,38^.

**Fig 1:**
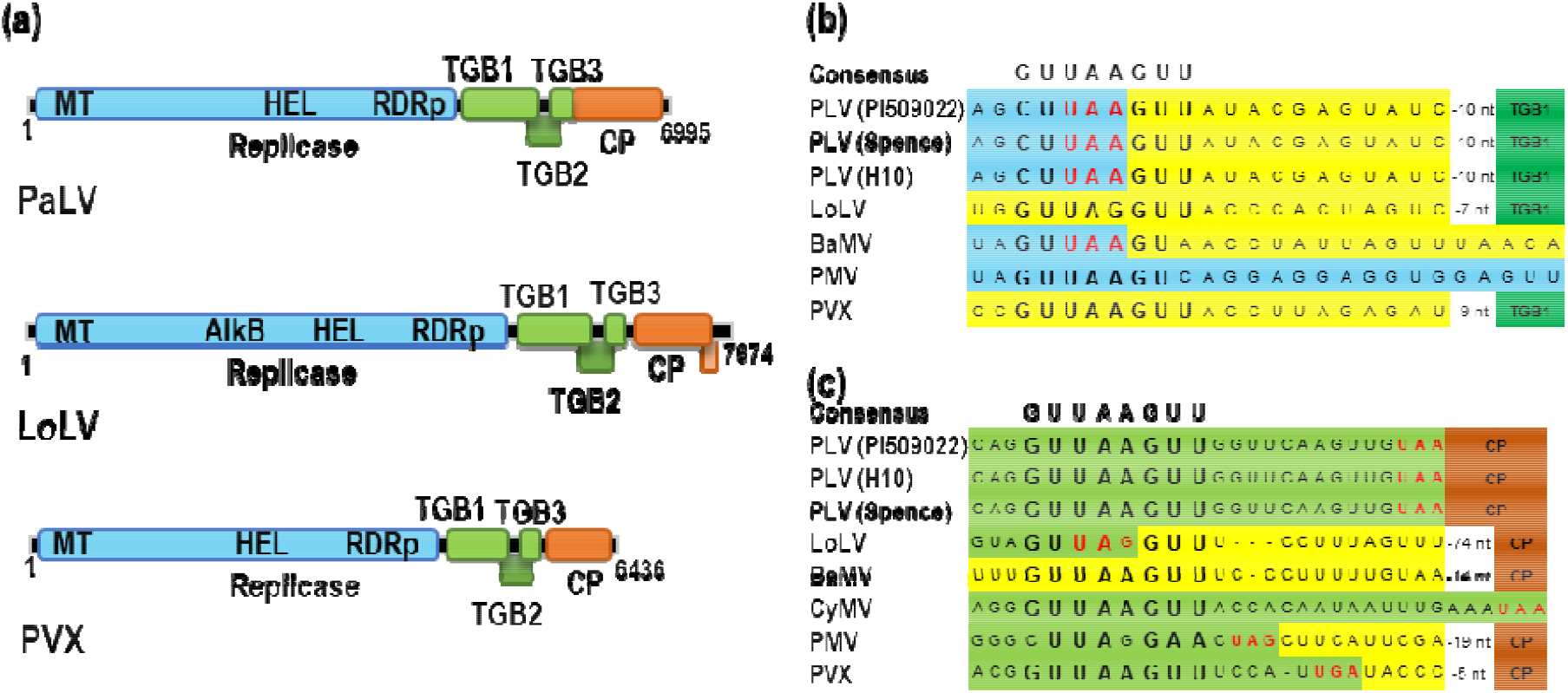
Diagram showing the genome organization of PaLV, LoLV, and PVX (a) and putative promoter sequences for sgRNAs of several viruses from genera, in *Lolavirus* and *Potexvirus* (b and c). (a) The five proteins enc ded by the viral genome are indicated and in LoLV, the extra small overlapping ORF with CP is also shown. (b and c) Alignment of the putative promoter sequences upstream of TGB1 (b) and CP (c). The consensus sequences at the top as reported in Kim & Hemenway, 1997, are indicated. Light blue and deep green color denotes RdRp and TGB1 encoding regions, respectively (b). Light green and brown color denote TGB3 and CP encoding regions, respectively (c). Yellow color indicates the intergenic regions and the three red nucleotides indicate the stop codons. BaMV: Bamboo mosaic virus (NC_001642); CyMV: Cymbidium mosaic virus (NC_001812); LoLV: Lolium latent virus (EU489641); PMV: Papaya mosaic virus (NC_001748); PaLV: Paspalum latent virus; PVX: Potato virus X (MT752888).

Downstream of the replicase, overlapping ORFs 2, 3, and 4 in different reading frames encode for putative triple gene block (TGB) proteins named TGB1, TGB2, and TGB3. The molecular weights of the predicted proteins encoded by ORFs 2, 3, and 4 are 30.4 kDa, 13.5 kDa, and 7.8 kDa, respectively. These proteins have been shown to play a crucial role in virus movement ^39^. Moreover, TGB1 has been implicated in viral translation and suppression of posttranscriptional gene silencing ^39^. A putative octanucleotide consensus promoter sequence for sgRNA1, CUUAAGUU, from the *Alphaflexiviridae* family ^40^ was identified 21 nt upstream of ORF2 (see Fig. 1B). sgRNA1 directs the expression of at least one ORF and, potentially, all ORFs related to TGB protein. Of particular interest is the observation that the promoter sequence of sgRNA1 of PaLV overlaps with the stop codon of ORF1, while in LoLV, the promoter sequence is found in the intergenic region between ORF1 and ORF2 (Fig. 1B). Kim & Hemenway (1997) reported that any mutation in the octanucleotide was highly detrimental to PVX sgRNA1 accumulation in protoplasts and eliminated infection in plants, thereby highlighting the importance of this sequence.

The ORF 5, located in the proximity of the 3′ end of the viral genome, encodes the coat protein (CP), which has a molecular weight of 35.4 kDa. Interestingly, the start codon of ORF 5 is in frame with and immediately follows the stop codon of ORF 4 (UUG <u>UAA</u> **AUG** GCA; the stop codon of ORF 4 is underlined, and the start codon of ORF 5 is in **bold**) (Fig. 1A). The absence of an intergenic region between ORF 4 and ORF 5 is atypical among the genera belonging to *Alphaflexiviridae*. However, in the case of Cymbidium mosaic virus (CyMV) in the *Potexvirus* genus, only two nucleotides are present between ORF 4 and ORF 5. Furthermore, the putative octanucleotide consensus promoter sequence for sgRNA2, GUUAAGUU, located 14 nt upstream of ORF 5 within the ORF 4, was also determined (Fig. 1C). Similarly to mutations in the sgRNA1 promoter sequence, any mutation in the putative sgRNA2 octanucleotide promoter sequence of PVX eliminated infection, indicating the significance of this sequence upstream of the CP ^40^. As with LoLV, another AUG start codon is in frame with the first start codon, 201 nt downstream, suggesting the possibility of two forms of CP (35.4 kDa and 28 kDa) getting incorporated in the virion particle. This is a typical characteristic of the *Lolavirus* genus that remains to be validated for PaLV. Contrary to LoLV, an ORF 6 that overlaps the 3′ end of ORF 5 was not identified in PaLV.

### Comparison of CP_PaLV_ with CP_LoLV_

Due to the extremely low amino acid sequence identity (26%) of ORF 5 with LoLV, we investigated the CP sequences in greater detail (Table 1). The protein localization prediction program, Plant-mSubP ^41^, indicated that the CP of PaLV (CP_PaLV_) exhibited a low probability of localizing in plastids (chloroplasts) and had an almost equal probability of localizing in the cytoplasm or peroxisomes. Conversely, the CPs of LoLV (CP_LoLV_) and PVX (CP_PVX_) demonstrated a high likelihood of localizing in plastids, a finding that aligns with the previous report on localization of CP in chloroplasts ^42,43^ (Fig. 2A). Subsequently, the protein structure of CP_PaLV_ and CP_LoLV_ was modeled to compare their structural relatedness. For this purpose, both amino acid sequences were subjected to Alphafold2 protein structure predictions (Fig. 2). The structure of CP_PaLV_ was predicted with a lower confidence level than that of CP_LoLV_ (Fig. 2). The structural modeling of both CPs revealed the presence of three domains: i) flexible N-terminal, ii) the core, and iii) C-terminal extensions. These domains are analogous to the crystal structures of CPs of PapMV and *Pepino mosaic virus* from the *Potexvirus* genus ^44,45^. The N- and C-terminal extensions play a crucial role in the polymerization of CP ^45^. The core and C-terminal regions of the CP play a crucial role in protein-protein and protein-RNA interactions and have been found to show the highest degree of structural conservation in *Alphaflexiviridae* ^45^. Consequently, a comparative analysis of the core and C-terminal domains (151-300 and 106-280 aa in CP_PaLV_ and CP_LoLV_, respectively) of both CPs was conducted. As predicted, a high TM score, indicative of high structural similarity, was obtained upon the superimposition of the core and C-terminal regions of both CPs, which consists of conserved α-helical secondary structures (Fig. 2). Collectively, these findings serve to further highlight the similarities and differences between CP regions from PaLV and LoLV.

**Fig 2:**
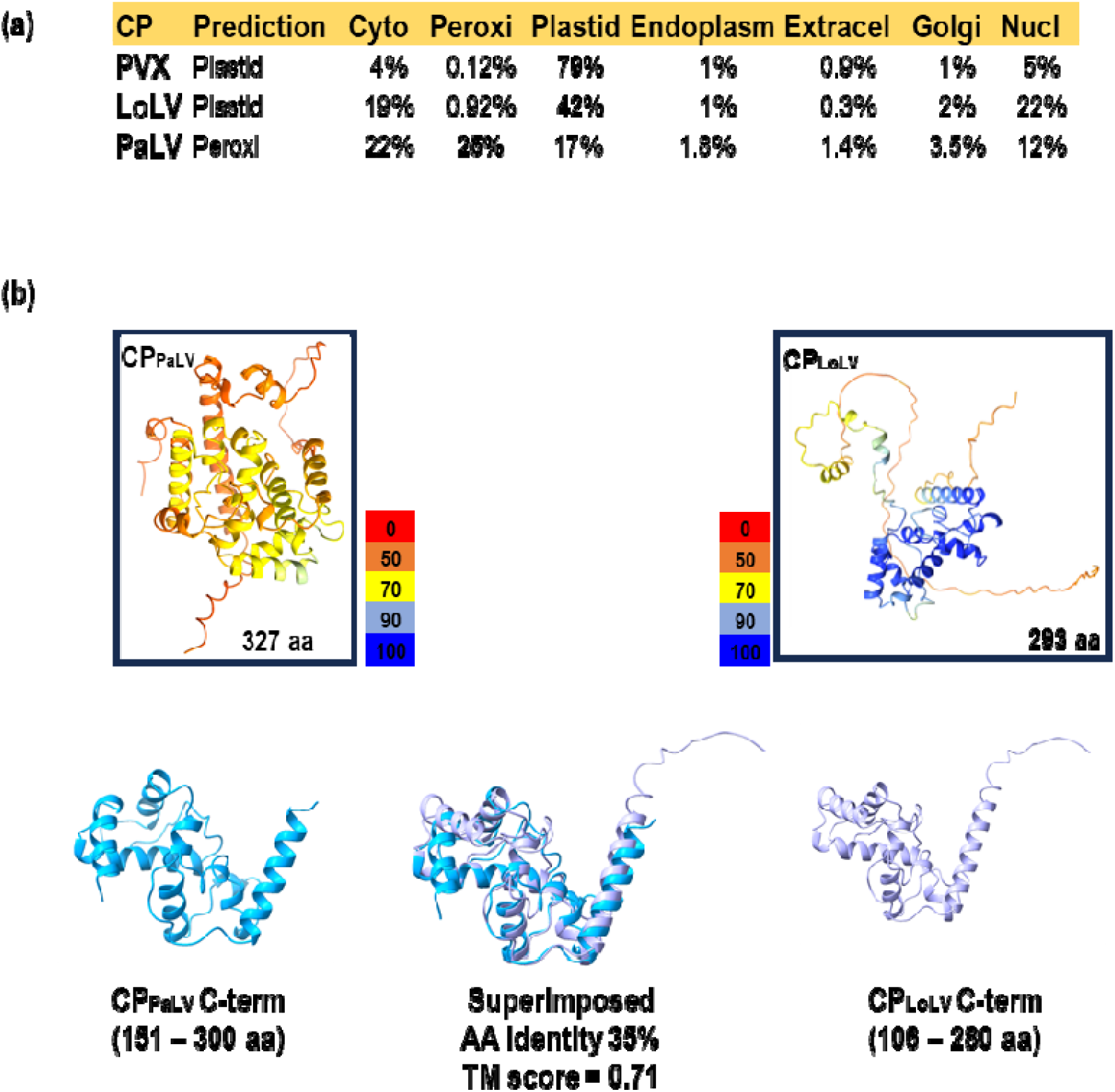
Comparison of CP_PaLV_ with CP_LoLV_. (a) A table showing the predicted localization of CP_PaLV_ and CP_LoLV_ in a plant cell as determined in a web server, Plant-mSubP. (b) Structure modeling of CP using Alphafold 2. Full-length structures of CP_PaLV_ and CP_LoLV_ are boxed. The numerical value shown at the bottom right corner indicates the number of aa in the CP. Color coding from red to blue indicates low to high structural confidence, respectively. The C-terminal of the CP_PaLV_ is superimposed on the corresponding CP_LoLV_ region with a high TM score.

### PaLV is a novel species in the genus *Lolavirus*

A phylogenetic tree was constructed to evaluate the evolutionary relationship of PaLV with other plant-infecting viruses from different genera of *Alphaflexiviridae*. The tree included 11 different species from various genera and the genome sequences of three PaLV isolates from this study. A putative viral sequence from *Alphaflexiviridae*, which is not a recognized species, obtained through data mining and thus submitted as Third-Party Annotation (TPA) was also included in the phylogenetic analysis (TPA_asm: Saltwater paspalum lolavirus 1 isolate Pas_vag, [SpLoV1; BK068268]). Overall, the tree showed four distinct clades with high bootstrap support for viruses from select genera in the family *Alphaflexiviridae*. The phylogenetic tree showed that PaLV isolates clustered with the genus *Lolavirus*, which contains only one established virus species, LoLV, with 100% bootstrap support. All PaLV isolates formed a sister branch to LoLV, indicating a unique cluster (Fig. 3). PaLV clade also contains the putative viral sequence, SpLoV1, which shares 97% nt identity with PaLV, suggesting that they belong to the same species. Interestingly, SpLoV1 was identified in the transcriptome shotgun assembly (TSA) database obtained from seashore paspalum deposited by Clemson University in the U.S., suggesting the presence of PaLV in the U.S. is consistent with our study ^46^.

**Fig 3:**
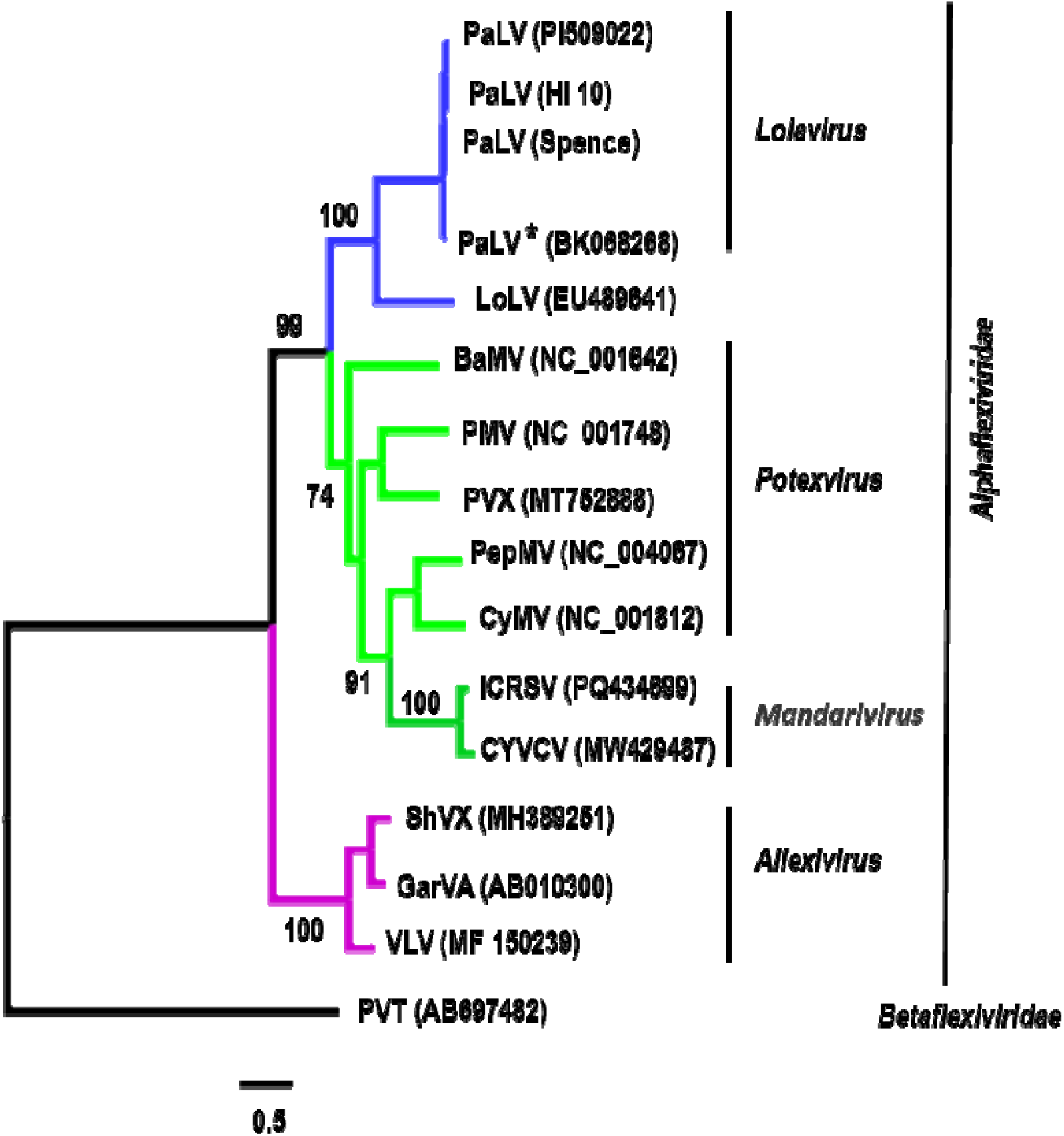
Maximum-likelihood phylogenetic tree based on aa sequences of replicase of plant infecting viruses from the *Alphaflexiviridae* family. The branch number indicates bootstrap support in percentage (out of 1000 replicates). The scale bar at the bottom denotes amino acid substitutions per site. The tree is rooted to an outgroup Potato virus T (PVT) from *Betaflexiviridae* family. * Indicate sequence obtained through data mining and is submitted in NCBI as Third party annotation assembly.

We also examined whether recombination between viral sequences played a role in the origin of PaLV; none of the algorithms implemented in RDP5 detected any signs of recombination. The phylogenetic data thus validates the ORF identity results, which indicates that PaLV shares the highest degree of identity with LoLV (Table 1). Based on the ICTV demarcation criteria in the family *Alphaflexiviridae*, new genera are established when the CP and replicase have less than 40% amino acid identity ^37^ However, the PaLV replicase showed 44% identity with LoLV, which is higher than the demarcation threshold for genera. However, PaLV meets the species demarcation criteria, which is less than 80% in either the CP or replicase (Table 1). Therefore, based on the ICTV guidelines, PaLV should be considered a new species and the second member of the *Lolavirus* genus.

### Genetic diversity of the PaLV population

A total of 11 CP sequences of PaLV isolates were obtained from different accessions of *P. vaginatum* and *P. distichum* (Supp Table 2). A phylogenetic tree at the nucleotide level of all eleven sequences revealed the presence of two distinct subgroups of PaLV isolates, designated clusters A and B (Fig. 4). Cluster A consists of all the isolates (except PI 509022) detected in *Paspalum* sp. originated in the US. In contrast, cluster B consists of all isolates (except K9) that originated from the rest of the world (ROW). This indicates a possible geographic-based separation with some level of population gene flow. For instance, Argentinian isolates from accessions PI 508737 and PI 509022 are observed in both sub-groups (Fig. 4 and Supp. Table 2). Gene flow among the PaLV population is possible, as supported by the *F*_*ST*_ value of 0.124. A *F*_*ST*_ of 0.124 indicated a low degree of spatial structure and genetic differences between the population ^47,48^. Because of the small number of accessions analyzed, the fact that all accessions were maintained in a single location for multiple years, and the geographic origins could possibly refer to sites where *Paspalum* plants were maintained before being donated to NPGS ^49^, whether the two PaLV populations indeed have different geographic origins and distributions will need to be confirmed. Nevertheless, similar observations of low to high genetic differences were also made in PVX populations, where isolates from different continents were analyzed ^47,48^.

**Fig 4:**
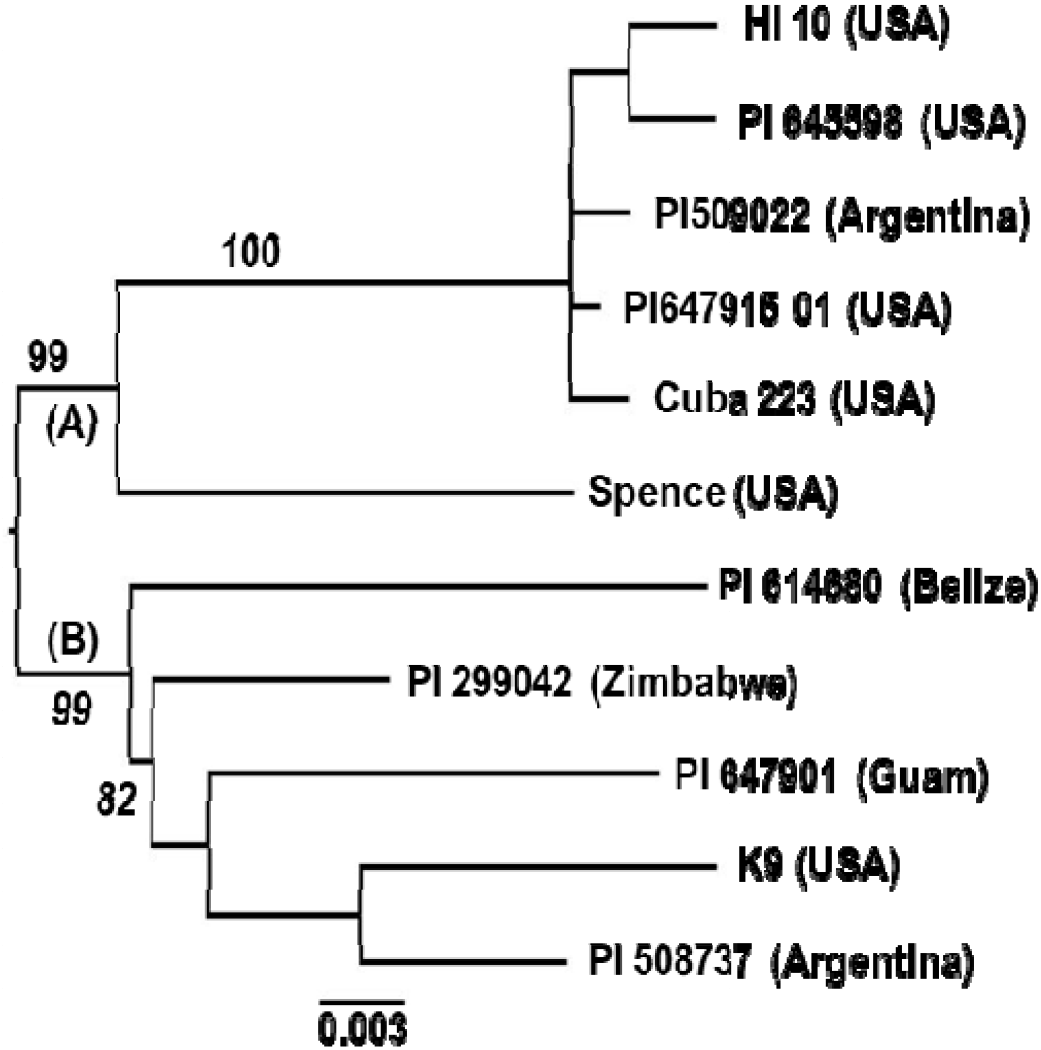
Maximum-likelihood phylogenetic tree based on the full CP nt sequences of PaLV. The branch number indicates bootstrap support in percentage (out of 1000 replicates). The scale bar at the bottom denotes amino acid substitutions per site. The tree is mid-point rooted. The country of origin is shown in parenthesis beside the accession numbers. Origin is not known for accessions PI647915 01 and K9.

To ascertain whether the moderate diversity could be attributed to selection, the dN/dS ratio was estimated, yielding a value of 0.042, suggesting the presence of strong purifying (negative) selection on CP. This finding is further corroborated by the observation of low genetic diversity (0.031 ± 0.004) of CP. These results are consistent with the expectation of CP sequence conservation given the numerous pivotal functions of CP within *Alphaflexiviridae*, including the activation of RNA translation ^50^, the facilitation of infection ^51^ and the encapsidation of viral genome RNA ^52^. Our findings are in agreement with studies on viral CP from different viral families that documented strong purifying selection pressure along with low genetic diversity ^47,48,53,54^. Altogether, it cannot be ruled out that seashore paspalums may have often been exchanged between countries due to their beneficial use in fields, leading to a low degree of spatial structure in the viral population instead of a very high spatial structure.

### Host Range Studies

Nine indicator plants representing nine plant genera were subjected to mechanical inoculation with leaf extract from PaLV-infected plants. All inoculated plants of the nine assayed genera were monitored over a period of 21 days post-inoculation (dpi) for symptom development. (Supp Table 1). All inoculated (virus and mock) plants were examined by RT-PCR. PaLV amplicons of the expected size were obtained with RT-PCR in: *Zea mays, Sorghum* spp., *Setaria italica*, and *Lolium multiflorum*. Interestingly, *T. aestivum* plants maintained at 16□ were also susceptible to PaLV infection based on RT-PCR. The infection of the aforementioned species indicates they are hosts of PaLV. On the contrary, no amplicons were amplified from the remaining inoculated species: *Hordeum vulgare, Avena sativa, Dactylis glomerata*, and *Miscanthus sacchariflorus*. None of the experimental hosts showed any symptoms of virus infection. Similarly, no symptoms were also observed in infected seashore paspalum germplasm. No symptoms were observed in mock-inoculated plants. In contrast, LoLV infections in *Lolium* plants were reported either to be asymptomatic or showing mild chlorotic streaking on the leaves ^21^ and in other plants systemic symptoms such as mosaic and vein netting showed at 15 dpi ^55^. This indicates that while PaLV shares similarity to LoLV, PaLV does not show severe or adverse effects due to the lack of virulent factors which can allow the virus to go undetected in some grasses.

### Screening of paspalum germplasm for PaLV infection

A one-step RT-PCR protocol was developed for screening PaLV infection. A primer pair was designed to target the RNA-dependent RNA polymerase domain in ORF1, amplifying a region of 605 bp. Thirty different *Paspalum* accessions maintained at UGA were analyzed. Two samples of butterfly bush were used as a negative control, and a PaLV-positive plant as a positive control. The RT-PCR results revealed the presence of PaLV infection in 27 out of 30 accessions (Supp Table 2), with an incidence of 90% in the tested *Paspalum* accessions. Notably, none of the negative controls showed the presence of PaLV, indicating the specificity of the primer pair. Given the global diversity of the *Paspalum* germplasm, it is intriguing to hypothesize that PaLV, like PVX, lacks the recent addition of AlkB domain in the replicase and is a virus that has been historically prevalent in *Paspalum* ^37–39,48^. Further studies are needed to confirm this hypothesis in historically curated *Paspalum* samples.

While other *Alphaflexiviridae* viruses have been easily recognized, PaLV, on the other hand, has not been detected immediately due to several factors. First, PaLV infection is latent, and symptoms are not expressed. Therefore, diagnostics are difficult without the use of molecular tools. While HTS can resolve this issue, routine detection is not practical with HTS. Second, *Paspalum* is not a commodity crop and garners less research attention. Nevertheless, in this study, we developed an RT-PCR assay for routine testing of PaLV that is available for turf research and diagnostics.

## Conclusions

In this study, we characterized a novel virus species, PaLV, belonging to the genus *Lolavirus* that was latently present in most *Paspalum* accessions maintained at UGA. Given that seashore paspalum is typically propagated vegetatively through sod, containerized material, stolons, and rhizomes, the virus can be easily disseminated during plant propagation ^56^. Here, we observed no discernible effect of PaLV infection on *Paspalum*. However, the asymptomatic nature of PaLV infections does not preclude the possibility of its impact on plant growth, breeding, and other plant physiological factors such as salt tolerance, a subject that requires further investigation. Furthermore, it is plausible that further adoption of *Paspalum* in the turf industry can potentially result in the emergence of more virulent PaLV strains due to vegetative propagation ^57–59^. Therefore, PaLV can potentially emerge as a challenge for turfgrass management with the lack of resistance traits in the *Paspalum* germplasm. Further studies should focus on screening *Paspalum* germplasm against PaLV through quantitative and semi-quantitative RT-PCR-based methods, similar to what was observed for resistance to fungal diseases in *Paspalum* ^7,60^. Nevertheless, even in the present context, it is imperative to implement control methods, such as using virus-free source materials to propagate paspalum.

## Data availability

## Acknowledgments

We gratefully acknowledge the funding for S.P. and K.D. from NSF award IOS-1915919.

## Author contributions

K.D., J.A.F., B.N.A. conceived and designed the research; S.B., T.F.S., conducted the experiments; A.L.P., T.F.S., A.H-W., S.P., C.D., collected and processed samples; S.B, X.H., P.A. analyzed the data from HTS; S.B., P.A., B.N.A. wrote the initial draft. All authors read, edited, and approved the manuscript

---

**Supplementary Table 1:** Bioassay to check for PaLV infectivity in different plants species.

| S.No | Crop | Bioassay Results |
| --- | --- | --- |
| **1** | ***Triticum aestivum* (16°C)** | **+** |
| **2** | ***Hordeum vulgare* (16°C)** | **-** |
| **3** | ***Triticum aestivum*** | **-** |
| **4** | ***Hordeum vulgare*** | **-** |
| **5** | ***Setaria italica*** | **+** |
| **6** | ***Sorghum spp*<sup>+</sup>** | **+** |
| **7** | ***Zea mays*** | **+** |
| **8** | ***Avena sativa*** | **-** |
| **9** | ***Lolium multiflorum*** | **+** |
| **10** | ***Dactylis glomerata*** | **-** |
| **11** | ***Miscanthus sacchariflorus*** | **-** |

+ presence of infection; - lack of infection

**Supplementary Table 2:** Screening of Paspalum Germplasm to check for PaLV infection.

| Sample | Name | Species <sup>(a)</sup> | From | PaLV PCR | Origin <sup>(a)</sup> |
| --- | --- | --- | --- | --- | --- |
| 1 | PI 576140 | *P. distichum* | UGA | + | Brazil |
| 2 | PI 647922 | *P. distichum* | UGA | - | Australia |
| 3 | PI 403999 | *P. distichum* | UGA | + | Australia |
| 4 | Durban | *P. vaginatum* | UGA | + | South Africa |
| 5 | PI 614679 01 | *P. vaginatum* | UGA | + | Bahamas |
| 6 | PI 299042 | *P. vaginatum* | UGA | + | Zimbabwe |
| 7 | PI 509018-3 | *P. vaginatum* | UGA | + | Argentina |
| 8 | K9 | *P. vaginatum* | UGA | + | Hawaii, USA |
| 9 | PI 647920 | *P. vaginatum* | UGA | + | Louisiana, USA |
| 10 | PI 647912 | *P. vaginatum* | UGA | + | Georgia, USA |
| 11 | PI 647918 | *P. vaginatum* | UGA | + | Florida, USA |
| 12 | PI 645598 | *P. vaginatum* | UGA | + | Texas, USA |
| 13 | PI 647916 | *P. vaginatum* | UGA | + | NK |
| 14 | Azul | *P. vaginatum* | UGA | - | NK |
| 15 | PI 614680 | *P. vaginatum* | UGA | + | Belize |
| 16 | PI 508737 | *P. vaginatum* | UGA | + | Argentina |
| 17 | PI647915 01 | *P. vaginatum* | UGA | + | Texas, USA |
| 18 | PI 647915 | *P. vaginatum* | UGA | + | Texas, USA |
| 19 | PI 509022 | *P. vaginatum* | UGA | + | Argentina |
| 20 | Cuba 223 | *P. vaginatum* | UGA | + | Hawaii, USA |
| 21 | Q36315 | *P. vaginatum* | UGA | + | Israel |
| 22 | PI 647908 01 | *P. vaginatum* | UGA | + | Georgia, USA |
| 23 | PI 377709 02 | *P. vaginatum* | UGA | + | South Africa |
| 24 | Kim 1 | *P. vaginatum* | UGA | + | Georgia, USA |
| 25 | PI 647901 | *P. vaginatum* | UGA | + | Guam |
| 26 | Hyb 5 | *P. vaginatum* | UGA | + | NA |
| 27 | Sea Dwarf | *P. vaginatum* | UGA | - | Florida, USA |
| 28 | KC 9 | *P. vaginatum* | UGA | + | Texas, USA |
| 29 | Spence | *P. distichum* <sup>(b)</sup> | UGA | + | Louisiana, USA |
| 30 | HI 10 | *P. vaginatum* <sup>(b)</sup> | UGA | + | Hawaii, USA |

(a) – The information on *Paspalum* species except for Spence and HI 10 and their origin were obtained from Eudy et al., 2017. NK and NA indicates not known and not applicable, respectively.
(b) – The information on *Paspalum* species were obtained from Spiekerman and Devos, 2020 + Positive PCR result; - Negative PCR result

